# Practical computational reproducibility in the life sciences

**DOI:** 10.1101/200683

**Authors:** Bjorn Gruning, John Chilton, Johannes Köster, Ryan Dale, Jeremy Goecks, Rolf Backofen, Anton Nekrutenko, James Taylor

## Abstract

Many areas of research suffer from poor reproducibility. This problem is particularly acute in computationally intensive domains where results rely on a series of complex methodological decisions that are not well captured by traditional publication approaches. Various guidelines have emerged for achieving reproducibility, but practical implementation of these practices remains difficult. This is because reproducing published computational analyses requires installing many software tools plus associated libraries, connecting tools together into the complete pipeline, and specifying parameters. Here we present a suite of recently emerged technologies which make computational reproducibility not just possible, but, finally, practical in both time and effort. By combining a system for building highly portable packages of bioinformatics software, containerization and virtualization technologies for isolating reusable execution environments for these packages, and an integrated workflow system that automatically orchestrates the composition of these packages for entire pipelines, an unprecedented level of computational reproducibility can be achieved.

---

Reproducible computational practices are critical to continuing progress within the life sciences. Reproducibility assures the high quality of published research by facilitating the review process that involves replication and validation of results by independent investigators. Further, reproducibility speeds up research progress by promoting reuse and repurposing of published analyses to different datasets or even to other disciplines. The importance of these benefits is clear, and vigorous discourse in the literature over the past several years^1–7^ has led to reproducibility guidelines at the level of individual journals as well as funding agencies.

However, achieving reproducibility on a practical, day-to-day level (and thus following these guidelines) still requires overcoming substantial technical challenges that are beyond the abilities of most life sciences researchers. There have been successful efforts aimed at addressing some of these challenges: Galaxy^8^, GenePattern^9^, Jupyter^10^, R Markdown^11^, and VisTrails^12^. These environments automatically record details of analyses as they progress and therefore implicitly make them reproducible. Yet they still fall short from achieving full reproducibility because they fail to preserve the full computing environment in which analyses have been performed. For example, consider an analysis executed on Galaxy, a Web-based scientific workbench used throughout the world. An analysis executed on a particular Galaxy server might include tools not found elsewhere, and therefore cannot be reproduced outside that server. Another example is a Jupyter notebook that includes tools specific to a particular platform and a distinct set of software libraries. There is absolutely no guarantee that such a notebook will produce the same results on a different computer. Here we introduce a solution that addresses all aspects of computational reproducibility by preserving the exact environment in which an analysis has been performed and enabling that environment to be recreated and used on other computing platforms.

While the need for reproducibility is clear and initial guidelines are beginning to emerge, research practices will not change until reproducible analysis becomes fast and automated. To make reproducibility practical, we have developed a three-layer technology stack composed of open, well-tested, and community supported components (Fig. 1). This three-layer design reflects steps necessary to make an analysis fully reproducible: (1) managing software dependencies, (2) isolating analyses from the idiosyncrasies of local computational environments, and (3) virtualizing entire analyses for complete portability and preservation against time.

**Fig. 1.**
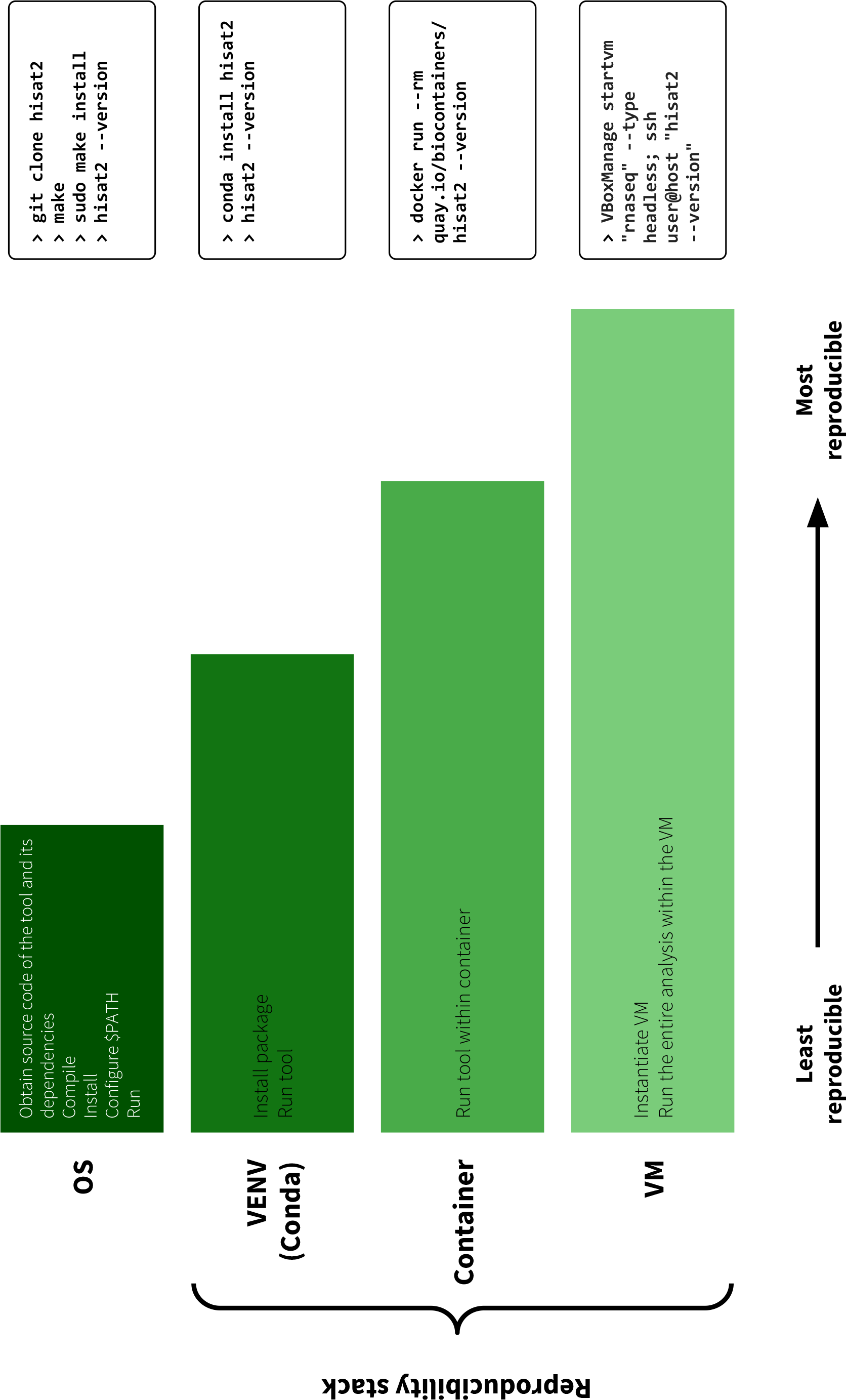
Figure 1. Software stack of interconnected technologies that enables complete computational reproducibility. It uses an example of the most basic RNA-seq analysis involving four tools. Our stack includes three components: (1) the cross-platform package manager Conda (https://conda.io) for installing analysis tools across operating systems, including virtualized environments that include all tools and dependencies at specified versions for performing a computational analysis; (2) lightweight software containers such as Docker or Singularity for using virtual environments and tool installations across different computing clusters both local and in the cloud; and (3) Hardware virtualization to achieve complete isolation and reproducibility. We have implemented this stack in the Galaxy scientific workbench (https://galaxyproject.org), enabling any Galaxy server to easily and automatically install all requirements for each Galaxy analysis workflow. This stack is also integrated into the Common Workflow Language (CWL; http://www.commonwl.org) reference implementation. Integration of or reproducibility stack into Galaxy and CWL demonstrates, for the first time, how analysis workflows can be shared, rerun, and reproduced across platforms with no manual setup. VENV = virtual environment, VM = virtual machine.

The first step, managing software dependencies, ensures that one can obtain the exact versions of all software used in a given analysis. Because most software tools rely on external libraries and analysis workflows use multiple tools, it is necessary to record versions of multiple tools and libraries. Given a multitude of operating systems and local configurations, ensuring the consistency of analysis software is a considerable challenge. Conda (https://conda.io), a powerful and robust open source package and environment manager, has been developed to address this issue. It is programming language and operating system independent, does not require administrative privileges, and provides isolated virtual execution environments. These features make Conda exceptionally well-suited for use on existing high-performance computing (HPC) environments as well as cloud infrastructure because precise control over the execution environment does not depend on system-level configuration or access. Conda explicitly supports installation of specific tool versions, even old ones, and allows the creation of “environments” where specific tool versions are installed and run. It is straightforward to create and maintain Conda package definitions, and this feature has led to rapid uptake of Conda by the scientific community. Leveraging Conda, Bioconda (https://Bioconda.github.io) is a community project dedicated to data analysis in life sciences that contains over 2,700 tool packages. Bioconda provides the top layer of our reproducibility stack. Bioconda packages are well maintained and include a testing system to ensure their quality. They are built in a minimal environment to allow maximum portability and are provided as compiled *binaries* which are archived, ensuring the exact executables used for an analysis can always be obtained. Conda environments and package management are agnostic to the underlying operating system. In contrast to other solutions such as Debian-Med^13^ or linuxbrew (http://linuxbrew.sh), Conda allows multiple versions of any software tool at the same time, provides isolated environments, and runs on all major Linux distributions, MacOS, and Windows.

While Conda and Bioconda provide an excellent solution for packaging software components and their dependencies, archiving them, and recreating analysis environments, they are still dependent on and can be influenced by the host computer system^14^. An additional level of isolation to solve this problem is provided by containerization platforms (or, simply, containers) such as Docker, Singularity, or rkt. Containers are run directly on the host operating system’s “kernel” but encapsulate every other aspect of the runtime environment, providing a level of isolation that is far beyond of what Conda environments can provide. From inside a container it is very difficult to access other containers or the host system itself. Containers are easy to create, which is a great strength of this technology. Yet it is also its Achilles heel, because the ways in which containers are created need to be trusted and, again, reproducible. This is why we generate containers automatically from Bioconda packages, and these automatically-created containers form the second layer of our reproducibility stack (Fig. 1). This has several advantages. First, container creation requires no user intervention, every container is created automatically and consistently using exactly the same process. A user of the container knows exactly what the container will include and how to use it. Second, this approach allows creation of large numbers of containers; in particular we automatically generate and archive a container for every Bioconda package. Third, this approach can easily target multiple container types. We currently build containers for Docker, rkt, and Singularity, and register them with Quay (https://quay.io). Since we build standard containers, other registries (e.g. DockerHub ^15^, BioShaDock^16^ or Dockstore^17^) can also be used. Because this container creation approach does not rely on the specification format of any particular system, additional container platforms and registries can easily be added as they become available. Finally, in addition to creating containers for single Bioconda packages, it is possible to automatically create containers for combinations of packages. This is useful when a step in an analysis workflow has multiple dependencies. Given any combination of packages with version, we can generate a uniquely named container which contains all of the required dependencies. When the combinations of dependencies required are known in advance these containers can be created automatically as well, for example we can create containers for all tool dependencies used in the Galaxy ToolShed (https://galaxyproject.org/toolshed)^18^.

Containers provide isolated and reproducible compute environments but still depend on the operating system kernel and underlying hardware. An even greater isolation can be achieved through virtualization, which runs analysis within an emulated virtual machine (VM) with precisely defined hardware specifications. Virtualization, which provides the third layer of our reproducibility stack (Fig. 1), can be achieved via commercial clouds, on public clouds such as Jetstream (https://jetstream-cloud.org), or by using virtual machine applications on a local computer (such as VirtualBox). While introducing this layer adds complexity and overhead, it provides maximal isolation, security, and resistance to time as emulated environments can be recreated in the future, regardless of whether the physical hardware still exists.

To make it easy for analysis environments and workflow engines to adopt this solution, we have implemented it in a Python library called *galaxy-lib.* Given a set of required software packages and versions, *galaxy-lib* provides utilities to either create a Conda environment with the required packages available, or to run the analysis with an appropriate container. Thus, a workflow can be executed in which every step of the analysis runs either using a dedicated conda environment or an isolated container. Support for this reproducibility stack has been integrated into the Galaxy platform, the Common Workflow Language reference implementation^19^ as well as Snakemake workflow engine^20^. Additional isolation and reproducibility can be achieved by running in a virtualized or cloud environment, for example using Galaxy CloudMan^21^ to run Galaxy on Amazon or Jetstream.

Reproducibility in computational life sciences is now truly possible. It is no longer a technological issue of “How do we achieve reproducibility?” Instead it is now an educational (or even sociological) issue of “How to make sure that the community uses *existing* practices?” In other words, how do we set a typical researcher (i.e., a graduate student or a post-doc) performing data analyses on the path of performing them reproducibly? While there are now several platforms that enable reproducibility, the technologies we describe here are both very general and easy to use. Thus, we offer the following recommendations:

1. **Carefully define a set of tools for a given analysis**. In many cases such as variant discovery, DNA/ Protein interactions assays, and transcriptome analyses, best practice tool sets have been established by consortia such as 1000 Genomes, ENCODE, and modENCODE. In other, less common cases, selection of appropriate tools must be done by consulting published studies, Q&A sites, and trusted, community supported blogs. There is no escape from this - methodological decision are as much a part of research as deciding on what cell lines to use or how to fine tune a qPCR assay. Because methodological decisions are vital to today’s biomedical research, it is essential to capture them so they can be shared with the scientific community. Recommendations 2-4 summarize our best practices for capturing and sharing these methodological decisions. To put this discussion on a practical footing, consider the simplest possible analysis of RNAseq data in which one needs to map reads against a genome, assemble transcripts, and estimate their abundance. The most basic set of tool for an analysis like this might include Trimmomatic^22^ to trim the reads, HISAT2^23^ to map the reads to a genome, StringTie^24^ to assemble and quantify transcripts, and Ballgown^25^ to perform differential expression testing.
II. **Use tools from the Bioconda registry and help it grow**. The registry at https://Bioconda.github.io provides the list of available packages. The three tools from our example are all available in Bioconda and can be used directly (Fig. 1). If a tool is not already available, one can either write a Bioconda recipe for the tool in question or request the tool to be wrapped by the Bioconda community by opening an issue at the project’s GitHub page. Note that using Bioconda-enabled tools is not just “good behavior for enabling reproducibility”. It is the easiest way to use these tools. First, it makes installation easy. Conda automatically obtains and installs all necessary dependencies, so the only requirement for installing, say, StringTie is opening a shell and typing “`conda install stringtie`”. Second, it makes analyses reproducible. Simply providing the output of “`conda env export`” with a manuscript allows anyone to easily obtain the exact version of the software used as well as *all* its dependencies.
III. **Adopt containers to guarantee consistency of results**. Analysis tools installed with Bioconda can be used directly. However the consistency of results (ability to guarantee that the same version of a tool gives exactly the same output every time it is run on a given input dataset) can be influenced by local computational environment. Because every Bioconda package is automatically packaged as a container the tools can be run from within the container in isolation providing a guarantee of result consistency. An example of this process for our RNA-seq example is shown in Fig. 1. As container technologies become widely available, we see this as the preferred way to use analysis tools in the majority of research scenarios.
IV. **Use virtualization to make analyses “resistant to time”**. Containers still depend on the host operating system, which will become outdated with time along with the hardware. To make an analysis “time proof” it is possible to use virtualization by encapsulating all tools, their dependencies, and operating system with a virtual machine image (VM). Virtualization has a higher barrier to entry and VMs alone have been criticized as being a “black box”. However virtualization enables time independence. Thus we recommend at least recording exactly the OS and hardware environment used for analysis so it can be recreated using VM technologies in the future. Executing analysis within a archived VM will always afford maximum reproducibility, and this will become easier as more compute resources move towards cloud-style operations.

In conclusion, we are reaching the point where not performing data analyses reproducibly becomes unjustifiable and inexcusable. Aside from hardening the software, the main challenges ahead are in education and outreach that will be critical for fostering the next generation of researchers. There are also substantial “cultural” differences among research fields in the degree of software openness that will need to be tackled. For example, genomics (which the authors represent and therefore are biased toward) has traditionally been quick in adopting new paradigms, while, for example, proteomics has been much slower^26^. We believe that this work is the first step toward making computational life sciences as robust as well established quantitative and engineering disciplines. After all - our health depends on it!

## Acknowledgements

The authors are grateful to Bioconda, BioContainers, and Galaxy communities as without these resources this work would not be possible. Nate Coraor provided critical advice on the project and edited the manuscript. This project was supported by NIH Grants U41 HG006620 and R01 AI134384-01 as well as NSF Grant 1661497 to JT, AN, and JG. RD was supported by the Intramural Program of the National Institute of Diabetes and Digestive and Kidney Diseases, National Institutes of Health. Additional funding is provided by German Federal Ministry of Education and Research (BMBF grants 031A538A & 031L0101C de.NBI-RBC & de.NBI-epi) to RB and BG.

